# Is Mycetoma a Vector-Borne Disease: The First Report on the Detection of *Madurella mycetomatis* in Ticks

**DOI:** 10.1101/775767

**Authors:** RS Azrag, SM Bakhiet, AM Almalik, AH Mohamed, AH Fahal

## Abstract

Currently, there is a massive gap the mycetoma knowledge in particular in its epidemiological characteristics, the infection route, the predisposing factors and the host susceptibility. With this background, the present cross-sectional descriptive entomological study was conducted to determine the possible role of arthropod vectors in the transmission of eumycetoma as well as the knowledge, attitude and practice (KAP) among the villagers towards that in a mycetoma endemic village at Sennar State, Sudan.

The study showed an abundance of indoors and outdoors arthropod vectors, and that included ticks, mosquitoes, sandflies, cockroaches and houseflies in the studied area. Ticks were more frequent, and they belonged to three genera and four species, and the later included *Hyalomma (H.) anatolicum* (11.03%), *Hyalomma (H.) rufipes* (0.67%), *Rhipicephalus (R.) everts* (73.1%) and *Amblyoma (A.) lepidium* (15.2%). The different types of the collected arthropod vectors were pooled in groups, and each group was screened for the presence of the *Madurella (M.) mycetomatis* DNA, the most frequent causative agents of eumycetoma in the studied area. The DNA was extracted, and amplification of the genomic rRNA genes was done by using universal pan fungal primers and specific *M. mycetomatis* primers. One pool containing *R. evertsi* DNA samples and one sample of *H. Rufipes* DNA gave positive results following PCR amplification of the universal fungal positive primers while *H. rufipes* sample gave positive results for *M. mycetomatis* using a specific primer. An association between the animals’ dungs, ticks and mycetoma transmission can be suggested from this study. However, further in-depth studies are needed to verify that.

**Author summary:** Mycetoma is a severely neglected tropical disease characterised by painless subcutaneous tumour-like swellings frequently noted in the extremities. There is a massive knowledge gap in transmission, infection route, and historically, it is believed to be associated with minor trauma caused by thorn pricks. This study was designed to determine the possible role of arthropods in mycetoma transmission in an endemic area in Sudan during the cold dry season. Pools of medically important arthropods were screened for mycetoma causative agents using DNA based method. The villagers’ habits and knowledge on arthropod vectors were examined using a pre-designed questionnaire. The results showed various presences of many arthropod vectors. Ticks were found in high prevalence, and densities in domestic animals found inside houses and the villagers had high contact level with the ticks in comparison to other vectors. The study reports for the first time, the detection of the causative agents of mycetoma in a pool of ticks. More studies on the possible role of ticks in the transmission of mycetoma diseases are badly needed to delineate the possible role of ticks on transmission of mycetoma.

## Introduction

Mycetoma is a chronic specific granulomatous progressive subcutaneous inflammatory disease, of both bacterial (actinomycetoma) and fungal (eumycetoma) origin [1]. The most common causative agents include the fungus *Madurella mycetomatis* and the actinomycetes *Nocardia brasiliensis, Actinomadura madurae, Streptomyces somaliensis*, and *Actinomadura pelletierii* [2]. Mycetoma is endemic in many tropical and subtropical regions, and Sudan is reported to have the highest incidence [3]. Most of the affected patients are of poor socio-economic status and low health education level and hence the late presentation and poor management outcome [4,5].

Being, a gravely neglected tropical disease, there is a massive knowledge gap in its pathogenesis and epidemiological characteristics. The latter include the disease susceptibility, resistance, transmission route and the incubation period, and that had led to difficulty in designing an objective control or preventive programme [6]. For the disease infection route of entry, the popular theory is, the infection is established following the traumatic subcutaneous inoculation of the causative agents following minor trauma caused by thorn pricks, sharp objectives, animals bites, and others [6,7]. However, in mycetoma endemic areas, the habit of going barefooted is frequent, and the minor injuries and thorn pricks are abundant. Thus it is expected that the incidence of mycetoma to be higher if this theory is true. Furthermore, many patients have no history of local trauma at the mycetoma site.

The primary reservoir of the causal agents is believed to be soil as several causative agents were cultured or their DNAs were isolated from the soil. That included *Actinomadura madurae, Actinomadura pelletierii, Nocardia asteroides, Nocardia brasiliensis, Streptomyces somaliensis* and among the fungi, *alciformispora senegalensis, Madurella mycetomatis, Neotestudina rosatii* and *Scedosporium boydii* [8–14]. Furthermore, it was observed that mycetoma endemic villages are characterised by poor hygiene and overcrowded houses and their proximity to the animals’ enclosures and their dungs. Also, the thorny trees and bushes are plenty; thus, several environmental factors are believed to predispose to mycetoma [15]. In mycetoma endemic villages with such poor environmental and hygienic conditions, it is expected to have rich collections of arthropod vectors able to transmit many diseases. With this background, this study was conducted to determine the role of arthropod vectors in an endemic village in Eastern Sennar, Sennar State, Sudan in the transmission of mycetoma. In this communication, we report for the first time a preliminary data on the possible role of arthropod vectors in the transmission of *Madurella mycetomatis* in the studied village.

## Materials and Methods

### Entomological surveillance

This descriptive cross-sectional study was conducted at Wad Al Nimar village, a mycetoma endemic village during the cold dry season. The village is located on the western bank of Aldindir River, Eastern Sennar locality, Sennar State, Sudan. It has a tropical climate with an annual rainfall of 600 mm and with varies relative humidity of 18% to 80% and temperature varies between 20^0^C and 40^0^C. The soil is mainly black cotton one, which cracks during the dry season and expands when the rain commences. The area has relatively rich natural vegetation with forest that cover about 34% of Sennar State ground. The State vegetation is characterised by savannah low rainfall trees and bushes dominated by *Acacia (A) seyal, A. Senegal, A. nilotica, A. melliferaand Balanites aegyptiaca*. Most buildings are made of mud that is mixed with animal dunglocally known as zebalah. Cracks on the walls of traditional houses made of zebalah provide suitable places for arthropod vectors to hide and rest. All houses include animal shelters for domestic animals. Aldindir River banks host many small farms of vegetables and fruits.

### Collection methods

#### Collection of indoor and outdoor disease vectors

Entomological surveillance for indoor resting and outdoor disease vectors was carried out in twenty randomly selected houses at the studied village. Informed written consent was obtained from the head of each house. In each of the selected houses, a combination of different collection methods was conducted, and that included light traps, sticky paper traps, active search and Knockdown methods[16]. Light and sticky paper traps in addition to active search methods were used for the collection of outdoor nocturnal vectors from animal shelters. Two light traps were suspended with the fan 40-50cm above the ground level. Traps were set one hour before the sunset and collected early next morning before the sunrise. The traps were transported to the field laboratory where sandflies and mosquitoes were sorted by sex and genus and were preserved in 70% ethanol for later identification to species level. For sticky trap collection, A4-sized white sticky traps (10 per night for three nights) coated with sesame oil were used to capture sandflies from outdoor habitats. A set of 10 sticky traps were hung vertically in a row of 30 cm above the ground supported by wooden sticks. Sticky traps were removed early in the morning, sandflies were removed using forceps and stored in 80% ethanol in labelled vials for further identification.

The Knockdown and active search collection (direct pick up) methods were used to collect indoor (inside rooms) diurnal resting mosquitoes and sandflies vectors. For Knockdown method, the rooms were sprayed early morning between 6:00 and 8:00am with commercial insecticide (Pif Paf). White sheets (2 x 2 meters) were laid on all flat surfaces over the entire floor and beds in the room, and all doors and windows were closed. Rooms were sprayed in a clockwise direction and care was taken to start spraying from the roof and all open spaces or holes in the walls until the room was filled with a fine mist. Then the room was quickly closed. After about 15 minutes, the door was opened, and the sheets were picked one at a time from their corners. The sheets were carried outside, and all knocked down arthropods were collected outside the rooms in daylight using forceps. Adult mosquitoes were stored dry in labelled Petri dishes using silica gel. Other arthropods were preserved in 80% ethanol in labelled falcon tubes.

Collection of tick samples from domestic animals was carried out by a veterinary doctor in addition to two trained volunteers from the local community. Ticks were directly collected from the invested domestic animals found in animal shelters (cows and goats only) and preserved in labelled Petri dishes using silica gel. Also, an active search for immature stages of ticks was carried out in some animal shelters. Search was focused on specific areas that included underneath fresh and dry animal dungs, water tanks and fresh plants offered as animal feed.

### Identification of arthropod Vectors

The morphological identification of the collected mosquitoes, sandflies and ticks was done according to keys used for the Afrotropical region [17–20]. Female mosquitoes and ticks were identified morphologically under a dissecting microscope. All collected sandflies (females and males) were sorted out as *Phlebotomus* or *Sergentomia* under a dissecting microscope then samples of *Phlebotomus* species were mounted on microscope slides in Puris media with their heads separated from thoraxes and abdomen. Identification of males was based on the morphology of external genitalia, and for females, identification was based on the pharynx, antennal features and spermathecae.

### Knowledge, attitude and practice (KAP) survey towards arthropod vectors

A KAP survey was conducted among the villagers in the studied village to determine knowledge, attitude and practice of villagers towards medically important arthropod vectors in the village. The survey was conducted by a team of two researchers and one health officer from the local community. A predesigned questionnaire was first tested in the field prior to the data collection for validation. The designed KAP questionnaire consisted of 49 questions which were grouped on 14 major sections. Eighty-one randomly selected villagers participated in the study after informed consent.

### Molecular screening of arthropod vectors

Pools of all collected mosquitoes, sandflies and tick samples were screened for the presence of *Madurella mycetomatis* using a modified method for the amplification of the (ITS1)-5.8S-ITS2 DNA region. Universal fungal and specific *Madurella mycetomatis* primers were used. The technique used was validated at the Mycetoma Research Center, Sudan.

### DNA extraction

DNA was extracted from pools of arthropod vectors. The DNA extraction was done using a modified protocol for DNA purification from tissues (QIAGEN KIDS). Fifteen metal beats were added in microcentrifuge tube. 500μl from ATL buffer was added, and the tube was put in the tissue lyser machine for 10mins/frequency 30, centrifuged at 10,000x for one minute. The supernatant was placed into a new tube, and 25μl proteinase K was added and incubated in a water bath at 56°^0^C for 30mins. Samples were centrifuged at 8000g for three mins. The clear supernatant was put into a new tube, and 300μl absolute ethanol was added and shaken briefly, then the tube was centrifuged to remove the drops from inside of the lid. The solution was put in a QIAGEN mini-column and centrifuged (8000g/3mins). 750μl AW1 solution was added into the column and centrifuged (8000g/3mins). The lid of the column was closed and was put in a new collection tube and centrifuged (15000g/1min). The column was placed in a new 1.5ml tube, and 70μl AE buffer was added inside the column and centrifuged (8000g/3mins). Samples were stored at −20^0^C.

### PCR using panfungal primers and *M. mycetomatis* species-specific primers

The isolated genomic DNA was amplified using PCR with panfungal primers ITS5 (5’-GGAAGTAAAAGTCGTAACAAGG −3’) and ITS4 (5’-TCCTCCGCTTATTGATATGC −3’) as described previously [23]. In the case of positive PCR with ITS4 and ITS5, *M. mycetomatis* species-specific primer was used 26.1A(AATGAGTTGGGCTTTAACGG); 28.3A (TCCCGGTAGTGTAGTGTCCCT). Reaction volumes of 20 μl contained 1 μl of genomic DNA, 1.25 U of AmpliTaq gold, 2 μl of 10× PCR buffer, 2 μl of 25 mM MgCl_2_, 2 μl of 2.5 mM deoxynucleoside triphosphate, and 1 μl of each 10 μM concentrated primers. The PCR products were amplified in an ICycler thermocycler (Aeris) set up with a first cycle of denaturation for 5 min at 95°C, followed by 40 cycles of denaturation at 94°C for 30 s, annealing at 56°C for 30 s, and elongation at 72°C for 30 s, with a final extension step of 10 min at 72°C. PCR products were visualised on 1% agarose gel after ethidium bromide staining.

### Statistical analysis

The data were managed by the Statistical Package for Social Sciences (SPSS) program (version 16). Descriptive statistics tests to measure the mean, percentages and ranges were used to analyse the obtained data on entomological surveillance and KAP study.

## Results

### Indoor resting and outdoor arthropod vectors

In this study, three groups of medically important arthropod vectors; mosquitoes, sandflies and ticks were collected from indoors and outdoors. For mosquitoes, forty-three indoor-resting mosquitoes of the *Anopheles*(*An*.) and *Culex*(*Cx*.) genera were collected and that included *Anopheles gambiae* complex (81.4%) and *Culex* species (18.6%). One hundred nineteen sandflies were collected; 73.1% belonged to *Sergentomyia* (*S*.) genus while 26.9% belonged to *Phlebotomus* (*P*.) genus. It was noticeable that cockroaches, houseflies and crickets were found in high densities indoors (Range of 15-30, 6-15 and 17-28 respectively). Other common non-arthropod vectors commonly found indoors included ants, silverfish and beetles.

Ticks were found in animal shelters present inside houses with very high levels of infestation and density; 100% infestation rate and densities that often range between hundreds and thousands in cows, sheep and goats. It is observed that unclean animal shelters, availability of animal dung, high density and diversity of animals accompanied with no veterinary services provided suitable environment for ticks. The contact level between humans, animals and ticks is considerably high inside animal shelters in comparison to grazing areas which led to a shift in the normal life cycle of ticks as breeding can take place inside animal shelters within the houses rather than at outdoor grazing areas.

Also, domestic animals (especially goats and sheep) were frequently found inside rooms and as villagers allow them to enter rooms during the hot hours in the middle of the day as the inside rooms provide a place with more suitable temperature. Also the same scenario happens during rainy season to protect their animals from rains.

One hundred forty-five ticks were randomly collected from animals (mainly goats and cows) found in animal shelters. The tick samples belonged to three genera and four species, and the later included *Hyalomma (H.) anatolicum* (11.03%), *Hyalomma (H.) rufipes* (0.67%), *Rhipicephalus (R.) evertsi* (73.1%) and *Amblyoma (A.) lepidium* (15.2%), (Table 1). Also, two nymphs of *Hyalomma* species were found during the active search underneath fresh cows dung in one of the animal shelters.

**Table 1:**
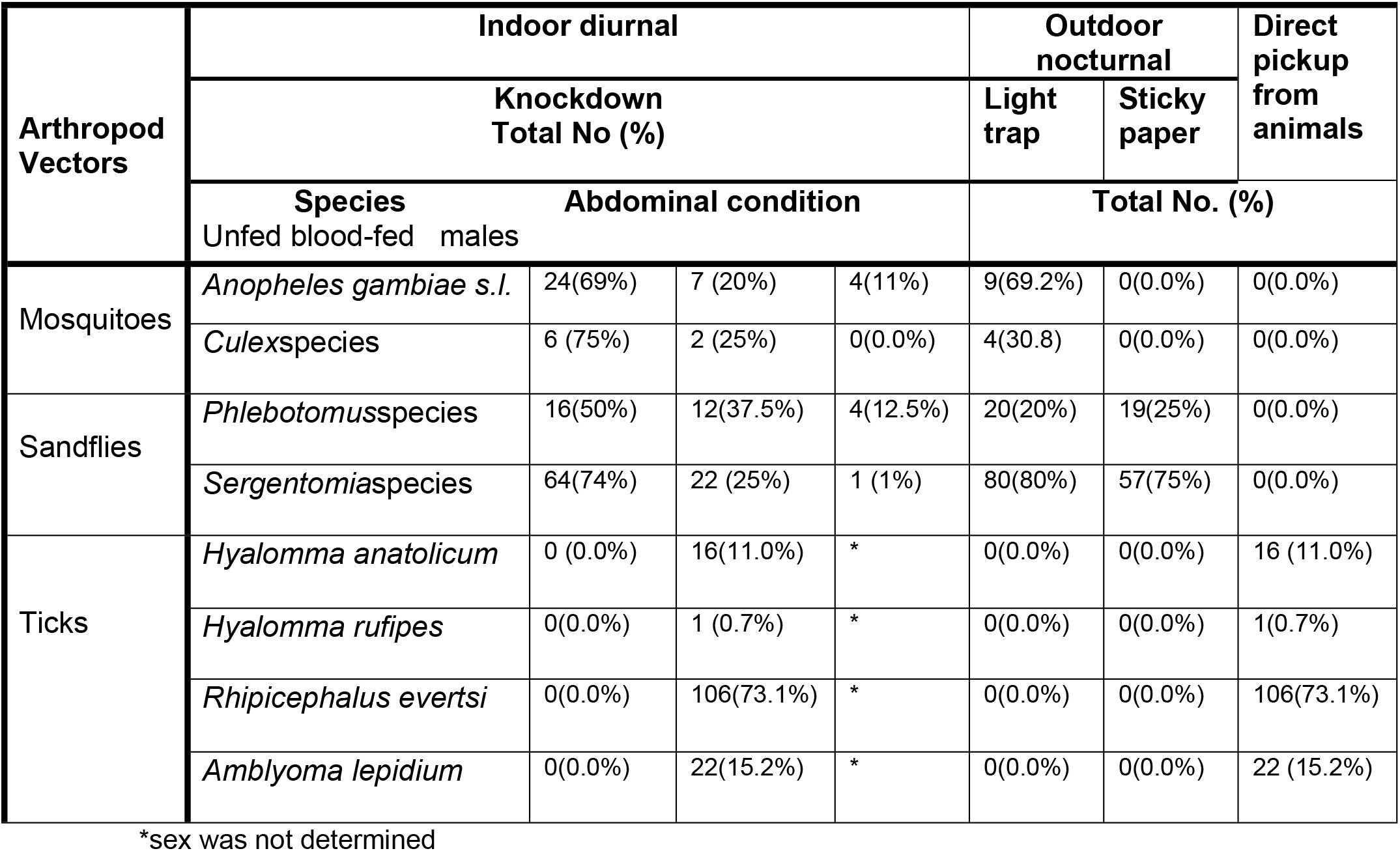
Indoor and outdoor arthropods collected during the cold dry season from the studied village.

### Results of KAP study

A total of 81 individuals (25 males and 56 females) were interviewed. Most of the males (72%) were farmers, and 89.2% of the females were housewives.26% of females reported a history of having a mycetoma infection at some stage. Both males and females had low educational level as 48% of males were illiterate, 24% had primary school education, 16% completed secondary school and 12% did not go to school and had some education at Khalwa (traditional place for learning Quran). For females, 53.5% had primary school education, 28.5% were illiterate, 14.2% completed secondary school and 3.5% studied at Khalwa. Villagers have a very simple life and main day time activities of the villagers included agricultural practice (92.5%), collection of firewood (61.7%) and grazing (24.6%). Most of the villagers (90.1%) reported an increased risk of animal bites during agricultural practices, 71.6% during the collection of firewood and 22.2% during grazing.

All villagers reported ticks as outdoor arthropod vectors, and 16% reported ticks as one of the arthropod species that bite outdoors while 7.3% reported ticks as one ofthe nocturnal arthropods. All villagers had very poor knowledge by diseases transmitted by arthropod vectors (especially ticks) and all of them reported that they do not use any protection measure to prevent them from arthropod bites (indoors and outdoors) (Table2).

**Table 2:** Results of the Knowledge, attitude and practice of the studied population toward local arthropod species in the studied village

| **Knowledge** |  |  |  |
| --- | --- | --- | --- |
| **Day time indoor arthropods** | **%** | **Day time indoor arthropods that bite** | **%** |
| Ants | 88.8 | Mosquitoes | 62.9 |
| Spiders | 50.6 | Ants | 70.3 |
| Crickets | 19.7 | Sandflies | 17.2 |
| Cockroaches | 38.2 |  |  |
| Flies | 16.0 |  |  |
| Mosquitoes | 07.4 |  |  |
| Fleas | 04.9 |  |  |
| **Day time outdoor arthropods** | **%** | **Daytime outdoor arthropods that bite** | **%** |
| Ticks | 100 | Ants | 100 |
| Flies | 44.4 | Mosquitoes | 72.8 |
| Ants | 100 | Ticks | 16 |
| Cockroaches | 20.9 | Fleas | 44.4 |
| Crickets | 08.6 |  |  |
| Fleas | 44.4 |  |  |
| Sandflies | 08.6 |  |  |
| **Indoor nocturnal arthropods** | **%** | **Indoor nocturnal arthropods that bite** | **%** |
| Mosquitoes | 100 | Mosquitoes | 90.1 |
| Crickets | 100 | Ant | 30.8 |
|  |  | Sandflies | 03.7 |
| **Outdoor nocturnal arthropods** | **%** | **Outdoor nocturnal arthropods that bite** | **%** |
| Mosquitoes | 91.3 | Mosquitoes | 100 |
| Crickets | 46.9 | Ants | 100 |
| Ticks | 07.3 |  |  |
| Cockroaches | 06.1 |  |  |
| **When do arthropods appear** | **%** | **Diseases transmitted by arthropod vectors** | **%** |
| Autumn | 76.5 | Malaria | 13.0 |
| During the year and increase in autumn | 23.4 | Leishmania | 02.0 |
|  |  | Other diseases | 0.00 |
| **Human Disease transmitted by ticks** |  |  |  |
| No | 100 |  |  |
| yes | 0.00 |  |  |
| **Attitude and Practice** |  |  |  |
| **Day time activities** | % | **Nighttime activities** | % |
| Agriculture | 92.5 | social activities | 92.5 |
| Collection of firewood | 61.7 | Watch TV | 06.1 |
| Grazing | 24.6 | Studying | 01.2 |
| Go to school | 06.1 |  |  |
| **Habits associated with arthropod bites** | % | **Protection measure against arthropod bites** | % |
| Agricultural practices | 90.1 | No | 100 |
| Collection of firewood | 71.6 | Yes | 00.0 |
| Grazing | 22.2 |  |  |

### *Madurella mycetomatis* molecular screening

Several arthropods pools were screened for the presence of *M. mycetomatis* DNA and that included one pool of 31 *Anopheles* females, one pool of eight *Culex* females and one pool of 28 *Phlebotomus* unfed females. Ticks samples were divided into pools according to the species and developmental stage. The *R. evertsi* samples were divided into five pools; one pool of 13 nymphs of larger size, one pool of 54 nymphs of medium size, one pool of 22 nymphs of small size, one pool of three adults with eggs and a pool of 14 adults. Other pools consisted of one pool of 22 *A. lepidium* adults, one pool of 16 *H. anatolicum* adults in addition to the one sample of *H. rufipes* adult.

The molecular screening showed that one pool containing *R. evertsi* DNA samples and one sample of *H. rufipes* DNA gave positive results following PCR amplification of the universal fungal positive primer while *H. rufipes* sample gave positive results for *M. mycetomatis* using a specific primer (Figs 1,2).

**Fig. 1:**
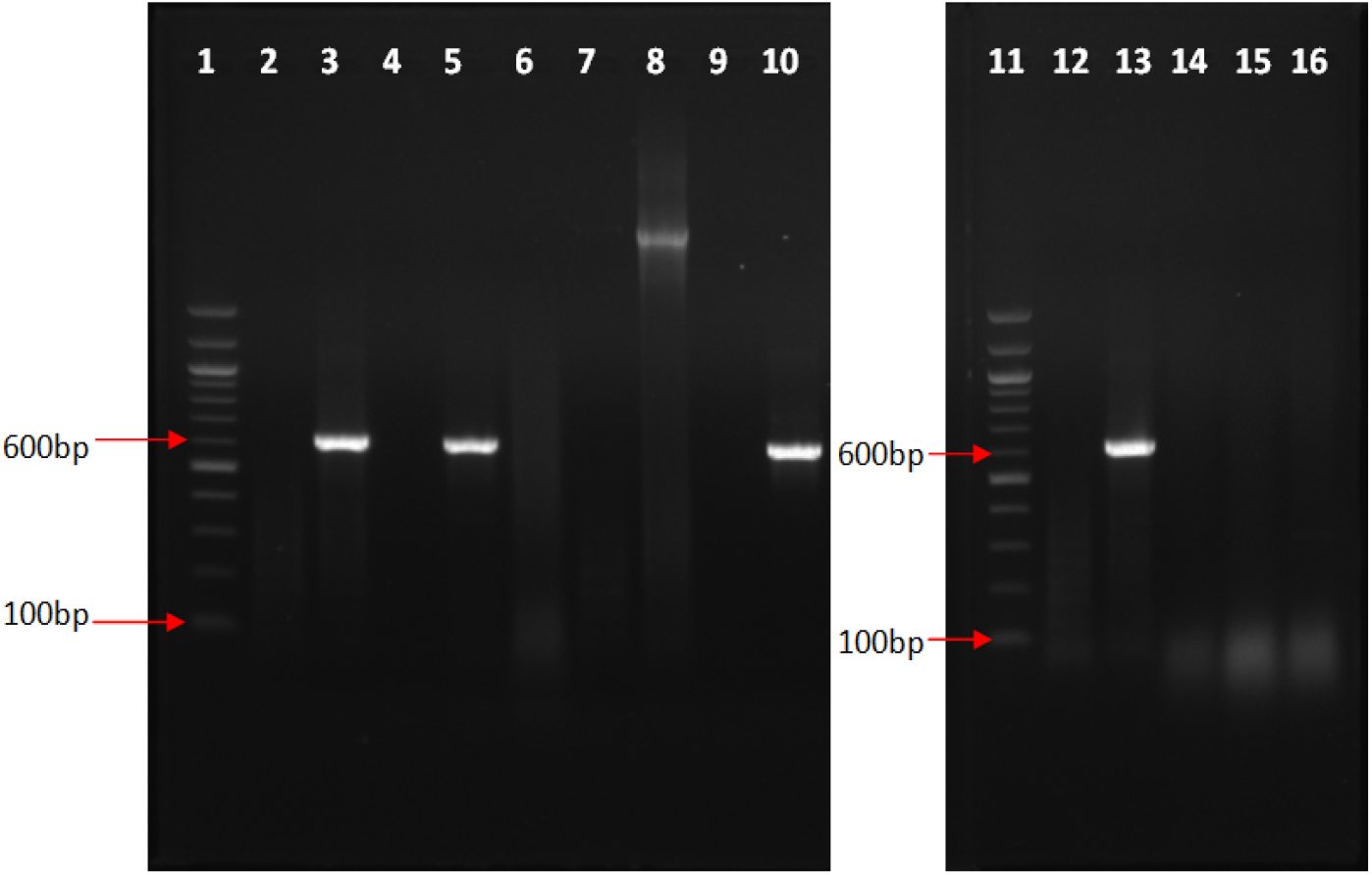
1% agarose gel visualising PCR products for the amplification of DNA pools using universal fungus primer. Lanes 1, 11: DNA ladder, lane 2: -ve control, Lane 3: +ve control, Lane 4: pool of R. evertsi, lane 5 = pool of *Rhipicephalus evertsi*, Lane 6: pool of *Amblyoma lepidium*, Lane 7: pool of *Rhipicephalusevertsi*, Lane 8: pool of *Hyalomma anatolicum*, Lane 9: pool of *Rhipicephalusevertsi*, Lane 10: *Hyalomma rufipessample*, Lane 12: -ve control, Lane 13: +ve control, Lane 14: pool of *Phlebotomus* sand flies, Lane 15: pool of *Culex mosquitoes*, Lane16: pool of *An. gambiae* complex.

**Fig.2:**
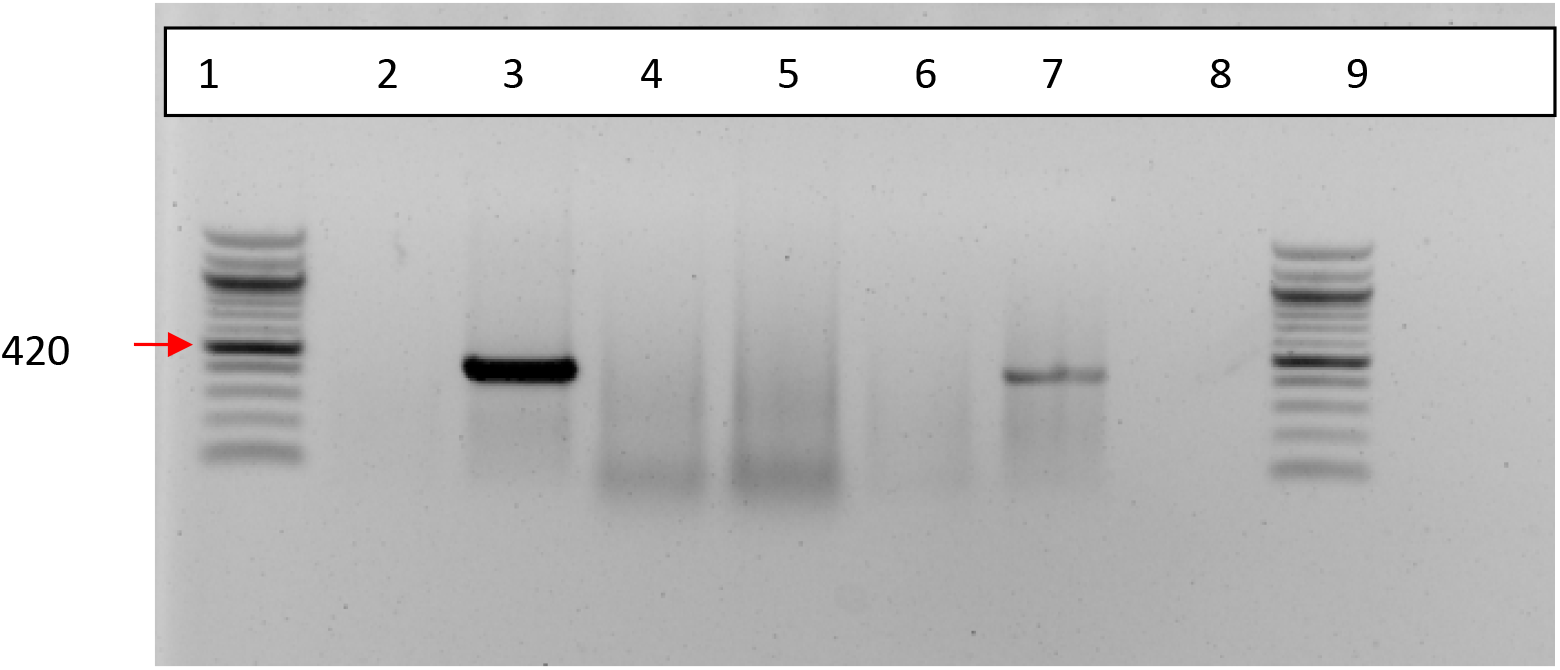
1% agarose gel visualising PCR products for the amplification of *M. mycetomatis* using specific primers. Lanes 1,9: 100 bp DNA Ladder, Lane 2: MM negative control, Lane 3: MM positive control, Lane 4: pool of *Rhipicephalus evertsi*, Lane 5: pool of *Hyalomma anatolicum, lane* 6: pool of *Amblyoma lepidium*, Lane 7: *Hyalomma rufipessample, Lane* 8: pool of *Rhipicephalus evertsi*

## Discussion

Presently, there are many controversies on the mycetoma causative organisms route of entry and disease susceptibility and resistance. However, mechanical implantation into the subcutaneous tissue is a prevalent theory [6]. Also, it is believed that certain environmental factors such as the poor hygiene, overcrowd housing, dirt, presence of animals’ dung, and others may contribute to this infection by providing a suitable environment for the causative organisms to survive but that is not clear yet [15]. Such poor environmental conditions also are suitable milieu for arthropod vectors to flourish. Considering all these, the current study was conducted to study the role of the common disease-transmitting arthropod vectors on mycetoma transmission in an endemic village at Sennar State, Sudan. The study reports, for the first time, the detection of *Madurella mycetomatis* DNA in ticks, and that may indicate their possible role in the transmission of mycetoma disease.

From the data obtained in this study, we can extrapolate a possible association between the ticks and mycetoma causative organisms’ transmission. Ticks are known as the most important vectors of many diseases affecting livestock and companion animals [21]. In addition, ticks are the second only to mosquitoes as vectors of human infectious diseases worldwide as they are known disease vectors for various diseases of protozoal, rickettsial, spirochetes, viral, fungal and bacterial origin and most of these diseases are of zoonotic origin. Since 1982, eight newly recognised tick-borne rickettsial; three species of ehrlichiae and three pathogenic species of the *B. burgdorferi* complex were reported to cause human diseases [22]. In Sudan, ticks and tick-borne diseases are wide spreading and cause substantial economic losses and constitute significant obstacles to the development of animal wealth. They are commonly causing important diseases such as tropical theileriosis, cowdriosis, babesiosis, anaplasmosis and avian spirochaetosis [23]. In Western Sudan, ticks are incriminated for the transmission of Crimean-Congo hemorrhagic fever in humans [24]. Tick species reported in this study were in agreement with the reported tick species in Sennar state, that are capable of producing animal diseases, specifically *Hyalomma anatolicum* species [25, 26]. Despite the economic and health importance of ticks it is believed that the reported knowledge on ticks and tick-borne diseases is still fragmentary and far from complete [23].

The present study showed high infestation of ticks accompanied with high densities inside animal shelters the studied village. In the ticks’ normal life cycle the drop off of the engorged females, oviposition and infestation happens outdoors (in grazing areas). However, in the studied village, high infestation rates and densities of ticks were due to unclean animal shelters, availability of animal dung, presence of high density and diversity of animals in addition to the high human contact level between animals and ticks points to a shift from the normal life cycle which completes indoors (in animal shelters) rather than in outdoors. Human contact level with ticks is higher in inside animal shelters in comparison to outdoor grazing areas.

The KAP study showed that villagers recognized ticks more than other indoor and outdoor arthropod. Also, men are in close and regular contact with immature stages of ticks during daytime indoor and outdoor activities, mainly during grazing and agricultural activities. Furthermore, women had high contact with ticks during the indoor activities in the animal shelters such as the milk milking process. In addition, both sexes also get exposed to ticks during the collection of wood and plants for fire and feeding animals from grazing areas. Moreover, villagers allow their animals to spend the daytime inside their rooms, especially during the hot dry season and the rainy seasons to protect them from the heat and rain. The KAP study showed that all villagers do not use any protection measure against arthropod bites, which points to the high exposure level to arthropod bites. We can postulate that villagers due to high direct contact with the ticks and the poor personal hygiene might get bitten more frequently by ticks, and the true percentage might exceed the percentage reported on this study. The reason is that ixodid ticks bites usually are usually painless and the immature ticks are often not detected in the human body due to their small size and hence the history of the local bite may not have recognised [22]. Moreover, got bitten by ticksis considered a social stigma therefore, probably some villagers are reluctant to report such event.

In the tropics, people often develop reactions to arthropods bites, and bacterial skin infections (pyodermias) usually follow such bites, stings and the mechanical trauma [27]. The type of reaction depends on the insect species, the age group and the human host reaction. The latter depends on the degree of previous exposure to the same or a related species of arthropod [28,29].

Ticks usually attach to human skin through their oral devices leading to diverse initial cutaneous manifestations, which can be classified into primary and secondary lesions. The primary one is caused by the attachment the tick to the host skin leading to severe skin inflammatory reaction due to the saliva anticoagulant substances and due to the penetration and permanence of the ticks’ mouthparts. The secondary lesions are due to the infections caused by rickettsia, bacteria, protozoa and fungi inoculated by the ticks [30]. From all these facts, we can then extrapolate that ticks mechanically can transmit mycetoma causative organism specifically under poor hygiene.

The life span of Ixodidae ticks ranges from several months to three years and they are less resistant to starvation and desiccation. Each ticks’ stages feed slowly by firmly attached to the host [22]. This indicates the ticks’ likelihood in diseases transmission, especially zoonotic diseases. Several studies showed that ticks prefer to bite lower extremities [22] and it is well known that the foot and hand are the most frequently affected sites (82%) in mycetoma affecting Sudanese patients [31] and that may support the ticks’ transmission postulation.

In Sudan, *M. mycetomatis* was isolated from soil and thorns samples and there is a possibility of a mycetoma-Acacia association [15, 32]. There is now evidence, from phylogenetic studies, that *Madurella* species are nested within the *Chaetomiaceae*, a family of fungi that mainly inhabit animal dung, enriched soil, and indoor environments [33]. In this study reported the collection of immature stages of ticks from underneath fresh cows dung inside animal shelters. This point to the possible association of ticks with the causative agents of mycetoma present on the soil and/or animal dung. The high contact level between animals, humans and ticks in animal shelters and the fact that ticks is frequently bite humans points to the possibility of incrimination of ticks on transmission of mycetoma disease.

The role of mosquitoes and sandflies on the transmission of mycetoma disease needs further investigations. The life cycle of immature mosquitoes is much associated with water. *Anopheles arabiensis*, a member of the *An. gambiae* complex is the main falciparum malaria vector in the eastern locality of Sennar State, and there is a marked seasonality on the transmission of malaria disease in this area [34–36]. However, in mycetoma, there is no clear seasonal variation, and hence, it is unlikely that these mosquitoes to have a role in the disease transmission.

Sandflies are the main vectors of visceral and cutaneous Leishmaniasis (CL) which are endemic in the studied area in eastern Sennar. In this area, visceral leishmaniasis is caused by *Leishmania donovani* and transmitted by *P. orientalis* while cutaneous leishmaniasis is caused by *Leishmania major* parasites and transmitted by *P. papatasi* sandflies [37, 38]. Transmission of leishmaniasis occurs most frequently outdoors with reports documenting indoor transmission [39–40]. The life cycle of sandflies is unknown until now. However, many reports showed that cracked cotton soil found in the study area could play a role as a resting or breeding places of adults vectors. Dogs, canines and the Egyptian mongoose, are possible reservoir host of visceral leishmaniasis in eastern Sudan [41]. However, the seasonality of leishmaniasis may not support the role of the sandflies in transmitting mycetoma.

In conclusion, an association between the animals’ dungs, ticks and mycetoma transmission can be suggested from this study. However, the role is unclear, but it can be postulated that tick bites cause minor injuries that may facilitate the inoculation of the mycetoma organisms into the subcutaneous tissue. However, this needs further studies. Furthermore, the role of the domestic animals as a possible mycetoma causative agents host reservoir and transmission needs meticulous investigation.

## Acknowledgements

The efforts of the Sennar State Ministry of Health staff in facilitating the study conduction were massive and enormous. Prof. Yassir Osman, Professor of Parasitology, Veterinary Research Center, Soba, Sudan, had kindly assisted in the ticks’ identification.

## Ethics Statement

The Mycetoma Research Centre, IRB approved the study.

